# Protection against experimental cryptococcosis elicited by Cationic Adjuvant Formulation 01-adjuvanted subunit vaccines

**DOI:** 10.1101/2024.04.24.591045

**Authors:** Ruiying Wang, Lorena V. N. Oliveira, Maureen M. Hester, Diana Carlson, Dennis Christensen, Charles A. Specht, Stuart M. Levitz

**Author notes:** Croda Pharma, Diplomvej 381, Lyngby 2800, Denmark (Dennis Christensen). Drs. Specht and Levitz share senior authorship. Corresponding Authors: Stuart M. Levitz, M.D., Department of Medicine, LRB317, UMass Chan Medical School, Worcester, MA 01605, USA; Charles A. Specht, Ph.D., Department of Medicine, LRB370D, UMass Chan Medical School, Worcester, MA 01605, USA.

## Abstract

The fungal infection, cryptococcosis, is responsible for >100,000 deaths annually. No licensed vaccines are available. We explored the efficacy and immune responses of subunit cryptococcal vaccines adjuvanted with Cationic Adjuvant Formulation 01 (CAF01). CAF01 promotes humoral and T helper (Th) 1 and Th17 immune responses and has been safely used in human vaccine trials. Four subcutaneous vaccines, each containing single recombinant *Cryptococcus neoformans* protein antigens, partially protected mice from experimental cryptococcosis. Protection increased, up to 100%, in mice that received bivalent and quadrivalent vaccine formulations. Vaccinated mice that received a pulmonary challenge with *C. neoformans* had an influx of leukocytes into the lung including robust numbers of polyfunctional CD4^+^ T cells which produced Interferon gamma (IFNγ), tumor necrosis factor alpha (TNFα), and interleukin (IL)-17 upon ex vivo antigenic stimulation. Cytokine-producing lung CD8^+^ T cells were also found, albeit in lesser numbers. A significant, durable IFNγ response was observed in the lungs, spleen, and blood. Moreover, IFNγ secretion following ex vivo stimulation directly correlated with fungal clearance in the lungs. Thus, we have developed multivalent cryptococcal vaccines which protect mice from experimental cryptococcosis using an adjuvant which has been safely tested in humans. These preclinical studies suggest a path towards human cryptococcal vaccine trials.

**Author summary:** Cryptococcosis is a fungal infection that poses great challenges to public health, especially in resource-limited regions with high HIV prevalence. Despite the urgent need, no licensed vaccines are currently available. In this study, we used a lethal mouse model of cryptococcosis to explore protection and immune responses elicited by vaccines consisting of recombinant cryptococcal proteins formulated with CAF01, an adjuvant that has an established safety and immunogenicity profile in human clinical vaccine trials. We discovered that while vaccines containing a single protein partially protected mouse strains, the protection was greatly augmented when the mice received vaccines formulated with multiple antigens. The lungs of vaccinated and infected mice had a robust influx of CD4^+^ T cells, many of which made the cytokines IFNγ and IL-17 when stimulated ex vivo. Moreover, we found the production of IFNγ directly correlated with clearance of fungi from the lungs. Cytotoxic CD8^+^ T cell responses were also observed, albeit in lesser numbers. Our promising findings from this preclinical research paves the way for future human cryptococcal vaccine trials.

## Introduction

Cryptococcosis is an opportunistic fungal infection caused by *Cryptococcus* species, mainly *Cryptococcus neoformans* and *C. gattii* [1]. Inhalation of airborne fungi into the lungs is thought to be the primary route of infection. While immunocompetent individuals generally clear or contain the infection, those with compromised immune systems are highly susceptible to clinically significant invasive disease or dissemination, commonly presenting as pneumonia, or, most severely, cryptococcal meningitis [2, 3]. The latter accounted for an estimated 152,000 HIV-associated cryptococcal meningitis cases in 2020, resulting in 112,000 deaths [4]. Even with antifungal therapy, morbidity and mortality rates remain high, particularly in resource-limited regions [5]. Thus, *C. neoformans* and *C. gattii* were categorized into the critical and medium groups, respectively, when the World Health Organization recently released their first Fungal Priority Pathogen List [6].

The development of vaccines represents a pivotal approach to prevent cryptococcal infections, yet no licensed vaccines are currently available [7]. Our laboratory focuses on identifying immunoreactive fungal proteins and developing effective vaccines. A promising strategy involves encapsulating these antigens within glucan particles (GPs) which are derived from *Saccharomyces cerevisiae* cell walls. GPs possess a 1,3-β-glucan outer shell, promoting receptor-mediated uptake by antigen presenting cells and enhancing immune responses to encapsulated antigens [8]. Among the twenty-three cryptococcal recombinant proteins tested in murine models, chitin deacetylase protein 1 and 2 (Cda1, Cda2), barwin-like domain protein 4 (Blp4), and carboxypeptidase 1 (Cpd1) emerged as highly promising protein antigens that afford significant survival advantages following pulmonary challenge with a highly virulent *C. neoformans* strain in both BALB/c and C57BL/6 mouse lines [9, 10]. Moreover, vaccination with these antigens encased in GPs elicits robust antigen-specific T helper (Th) 1 and Th17 responses critical for vaccine-induced protection [11].

However, challenges remain in formulating GP-subunit vaccines for large-scale production and translating these findings into clinical applications as GP-based vaccines have not been tested in clinical trials. To maximize our chances of developing a human vaccine, we have been evaluating adjuvants which have been successfully tested in clinical vaccination trials and which stimulate strong Th1 and Th17 responses when formulated with protein antigens. Notably, Cationic Adjuvant Formulation 01 (CAF01), a fully synthetic liposome-based adjuvant composed of the cationic quaternary ammonium salt N,N’-dimethyl-N,N’-dioctadecylammonium (DDA) and the glycolipid mycobacterial immunomodulator α,α′-trehalose 6,6′-dibehenate (TDB), emerged as a compelling adjuvant to pursue [12]. Vaccines are formulated simply by mixing antigens with CAF01, enabling large-scale, low-cost production. Similar to GPs, the CAF01 adjuvant works through caspase recruitment domain family member 9 (CARD9) pathways [13]. GPs are recognized by Dectin-1, while TDB contained in CAF01 is a Mincle ligand [14, 15]. By activating antigen-presenting cells (APCs) through a Mincle/CARD9 pathway, CAF01 induces a mixed Th1/Th17 response to formulated antigens [16]. Moreover, CAF01 has demonstrated safety and immunogenicity in human clinical trials of tuberculosis, chlamydia, HIV, and malaria vaccines [17-20]. Taken together, these studies lend a strong rationale for testing CAF01 as an adjuvant in cryptococcal vaccines. Thus, in the present study, we evaluated the protective efficacy of CAF01-adjuvanted subunit vaccines in a murine model of pulmonary cryptococcal infection and examined the mechanisms underlying vaccine-induced immunity.

## Results

### Efficacy of CAF01-adjuvanted monovalent subunit vaccines against lethal pulmonary infection in mice

In the first set of experiments, we investigated the protective efficacy of CAF01-adjuvanted monovalent subunit vaccines in a murine model of cryptococcosis using the highly virulent *C. neoformans* strain, KN99. BALB/c and C57BL/6 mice received three biweekly subcutaneous monovalent vaccinations formulated with the adjuvant CAF01. The cryptococcal protein antigens studied were Cda1, Cda2, Blp4, and Cpd1Δ. Two weeks after the final booster, the mice were subjected to orotracheal challenge with *C. neoformans* (Fig 1A).

**Fig 1.**
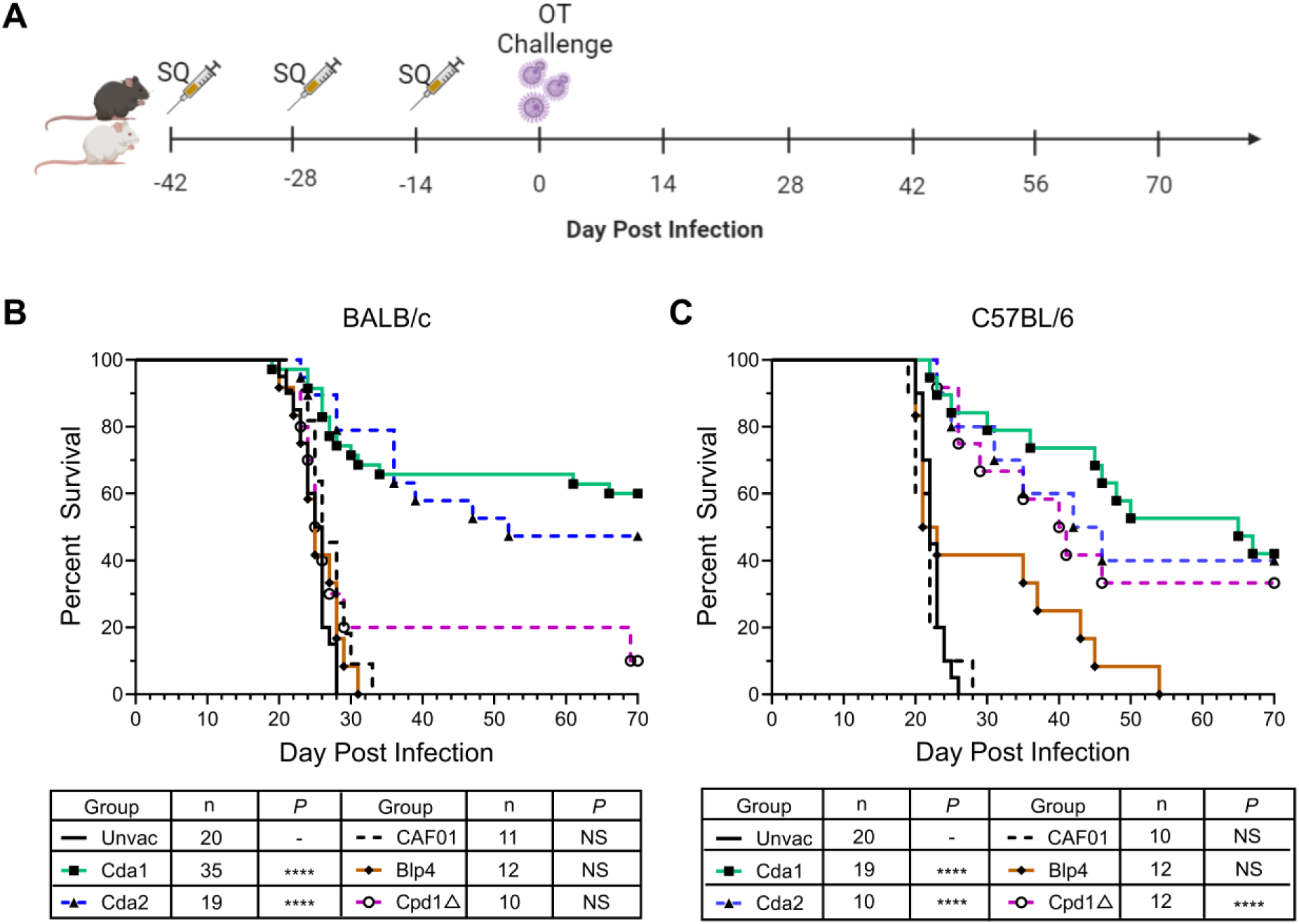
Protection induced by CAF01-adjuvanted monovalent subunit vaccine. (A) Schematic of vaccination strategy. WT BALB/c and C57BL/6 mice received a prime and two biweekly boosts with the indicated CAF01-adjuvanted vaccines. Mice were challenged with *C. neoformans* two weeks after the last boost, and then monitored for 70d for survival. The figure was created with BioRender.com. (B) Survival curves of BALB/c mice. (C) Survival curves of C57BL/6 mice. SQ, subcutaneous. OT, orotracheal. Unvac, unvaccinated mice. CAF01, mice receiving CAF01 without antigens. n denotes the total number of mice in the experimental group; each group included at least 2 independent experiments. ****, *P* < 0.0001; NS, *P* > 0.05, comparing the experimental group with Unvac mice.

For both mouse strains, unvaccinated mice and mice that received CAF01 alone succumbed to infection by 33 days post-infection (DPI) (Fig 1B, Fig 1C). For the mice vaccinated with a cryptococcal protein antigen in CAF01, protection varied as a function of the antigen and mouse strain. For BALB/c mice, Cda1 and Cda2 vaccination demonstrated statistically significant protection, with 60% and 47% survival at 70 DPI, respectively. A fraction of the mice vaccinated with Cpd1Δ survived to 70 DPI, however this trend was nonsignificant. No survival benefit was observed for the Blp4 subunit vaccine in BALB/c (Fig 1B).

For the C57BL/6 mice, significantly increased survival was observed with CAF01-adjuvanted vaccines containing Cda1, Cda2, and Cpd1Δ vaccines; survival varied from 33-42% survival at 70 DPI. Notably, all C57BL/6 mice succumbed following Blp4 vaccination and challenge, although an extension of survival was observed in 40% of the mice (Fig 1C). For both BALB/c and C57BL/6 mice, vaccination sites were inspected daily; no inflammatory reactions were observed.

### Enhanced protection against pulmonary cryptococcosis with CAF01-adjuvanted bivalent and quadrivalent subunit vaccines

The vaccine studies using single antigens adjuvanted in CAF01 demonstrated some of the antigens afforded protection against cryptococcosis in a murine model. However, protection, while significant, was only partial. We hypothesized that more robust protection would be obtained using combinations of antigens. Therefore, we investigated the efficacy of bivalent (Cda1+Cda2, referred as 2-Ag) and quadrivalent (Cda1+Cda2+Blp4+Cpd1Δ, referred as 4-Ag) subunit vaccines in BALB/c mice, following the vaccination and challenge schedule detailed in Fig 1A. These antigen combinations were chosen based on the experiments shown in Fig 1 and published studies with GP-adjuvanted subunit cryptococcal vaccines [10, 11]. After challenge with *C. neoformans*, weight and survival of the mice were monitored until a mortality endpoint was reached. Mice surviving to 70 DPI were euthanized.

Notably, the combination vaccination of 2-Ag protected 70% of BALB/c mice from subsequent lethal pulmonary infection, while 100% protection was achieved with the 4-Ag combination vaccination (Fig 2A). This degree of protection was greatly enhanced compared to single antigen vaccination. By 30 DPI, the unvaccinated mice exhibited 100% mortality. Moreover, unvaccinated mice exhibited minimal weight loss until 14 days post-infection, after which point they experienced a rapid decline in body weight, accompanied by signs such as scruffy fur, hunched back, rapid breathing, and reduced mobility (Fig 2B). In contrast, mice vaccinated with either the 2-Ag or 4-Ag combination vaccines began losing weight within the first few DPI, presumably due to the recall inflammatory response following infection (Fig 2B). The weights reached a nadir around 10 DPI, with an average weight loss of about 12% in both vaccinated groups. Subsequently, there was a steady recovery in body weight until the conclusion of the study.

**Fig 2.**
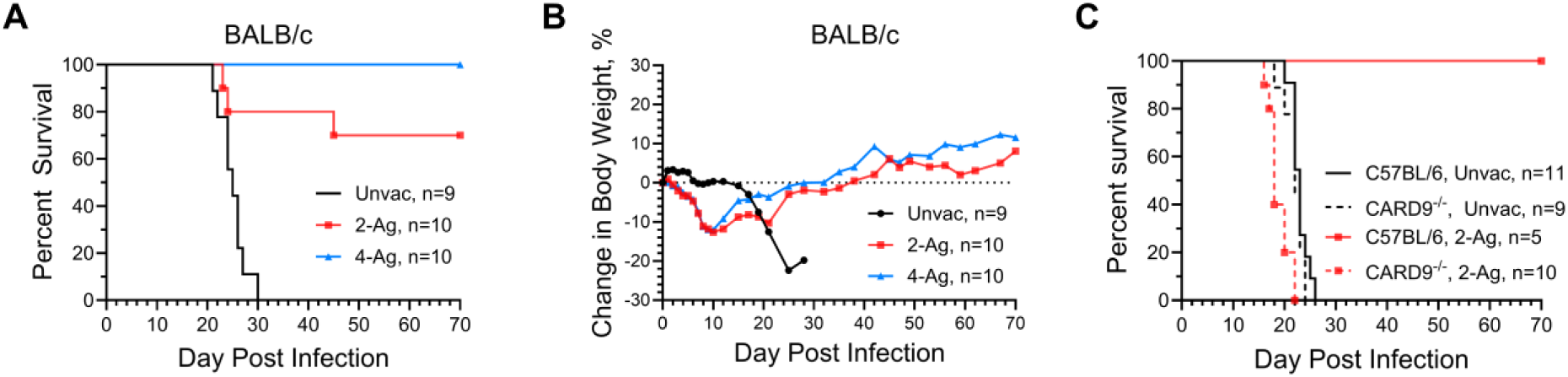
CAF01-adjuvanted combination vaccines enhance the protection of mice against experimental cryptococcosis. WT BALB/c mice received three SQ vaccinations with either 2-Ag or 4-Ag combination vaccines, following by orotracheal challenge with *C. neoformans*. Mice were monitored until death or study termination at 70 DPI for survival (A) and change in body weight (B). Data are from two independent experiments, each with 5 mice per group. Significant survival compared with unvaccinated mice was observed for both vaccinated groups in panel A (2-Ag, *P* = 0.0009; 4-Ag, *P* <0.0001). WT C57BL/6 mice or CARD9^-/-^ mice on the C57BL/6 background received 2-Ag vaccines following the same schedule. Mice were then monitored until 70 DPI for survival (C). Significant survival was observed in panel C, when comparing survival of vaccinated WT C57BL/6 and CARD9^-/-^ mice (*P* = 0.0006), or comparing vaccinated and unvaccinated WT C57BL/6 mice (*P* = 0.0003). Unvac, unvaccinated. 2-Ag, vaccinated with CAF01-Cda1+Cda2. 4-Ag, vaccinated with CAF01-Cda1+Cda2+Blp4+Cpd1Δ.

Furthermore, we assessed the 2-Ag combination vaccination in both wild type (WT) C57BL/6 and CARD9^-/-^ mice (Fig 2C). All of the unvaccinated mice succumbed to infection by 26 DPI. Meanwhile, all of the vaccinated C57BL/6 mice were protected to 70 DPI, at which time the experiment was terminated. Conversely, CARD9^-/-^ mice showed a complete loss of protection, consistent with CAF01 being a Mincle agonist, and Mincle’s utilization of CARD9 as its adaptor protein for downstream signaling [15].

### CAF01-adjunvanted subunit vaccines induce improved fungal control and augmented CD4^+^ T cell recruitment following cryptococcal infection

The above studies demonstrated CAF01-adjuvanted combination vaccines afforded up to 100% protection following cryptococcal challenge. In the next set of experiments, we investigated lung weights, colony forming units (CFUs), and immune responses in vaccinated and/or infected BALB/c mice at specific time points. Mice were either left unvaccinated or vaccinated with 2-Ag or 4-Ag vaccines. Within each cohort, mice were euthanized at specific time points for organ analysis: (1) 0 DPI, reflecting uninfected mice euthanized two weeks after the last vaccine boost; (2) 10 DPI, representing mice infected with *C. neoformans* and euthanized 10 DPI; and (3) 70 DPI, signifying mice infected with *C. neoformans* and euthanized 70 DPI. Note that the 70-DPI group did not include unvaccinated mice as they all succumbed to infection before 30 DPI.

Lungs from individual mice were collected and analyzed. As an indicator encompassing both infection and inflammation, lung wet weights were measured (Fig 3A). Lung weights were comparable between unvaccinated and vaccinated groups before infection. Ten days post-challenge, lung weights across all groups increased, with 4-Ag vaccinated mice exhibiting significantly heavier lungs than their unvaccinated counterparts. By 70 DPI, the lung weights in the 4-Ag vaccinated survivors had decreased but were still significantly higher than vaccinated uninfected lung weights at 0 DPI. There was a trend, albeit not statistically significant, towards greater lung weights in 2-Ag vaccinated mice at 10 DPI when compared with unvaccinated infected lungs. Moreover, at 70 DPI, the lung weights of 2-Ag vaccinated survivors declined to levels similar to what was observed before infection. No significant differences were observed between the 2-Ag and 4-Ag groups at any time point.

**Fig 3.**
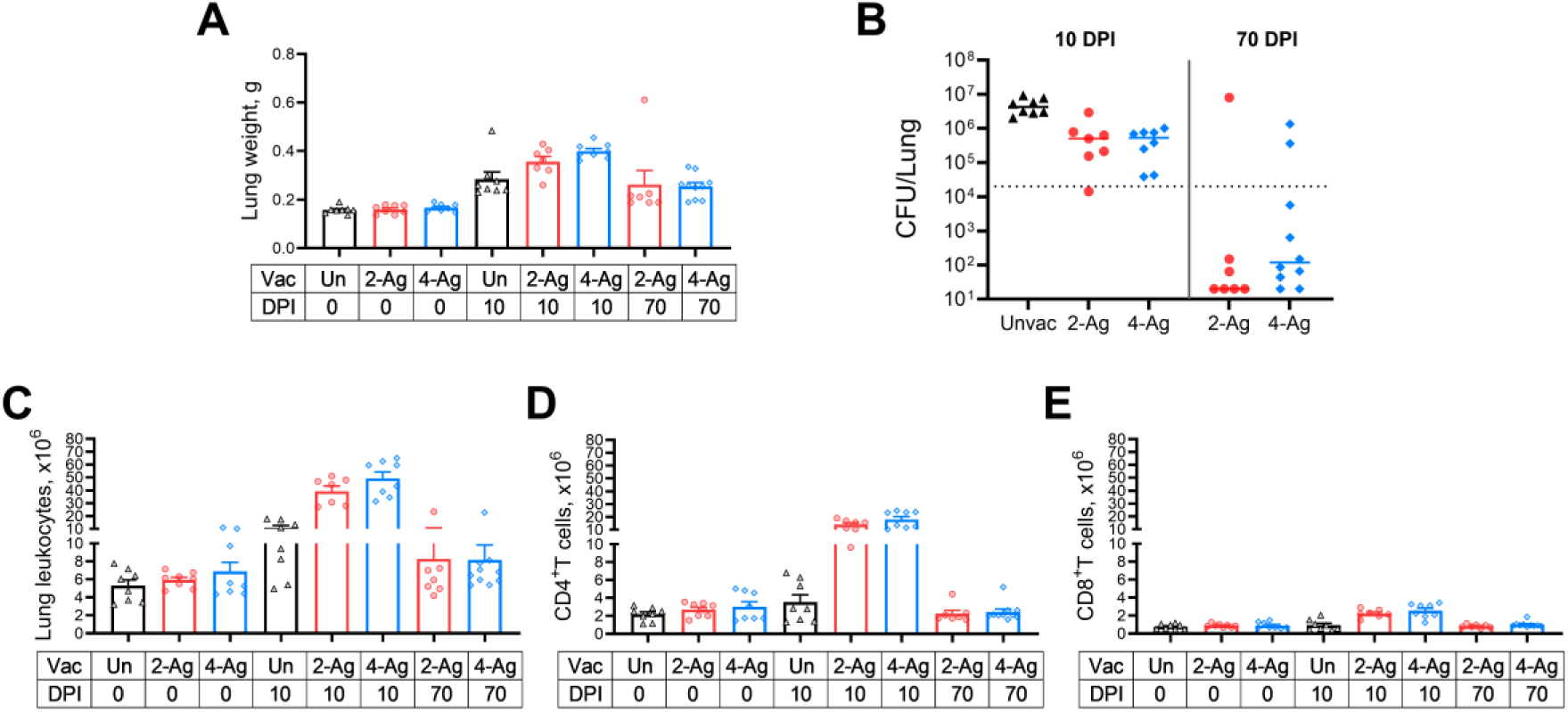
Fungal clearance and immune cell recruitment following CAF01-adjuvanted combination vaccines and/or infection. BALB/c mice received three biweekly vaccines with either 2-Ag or 4-Ag combination vaccines, followed by pulmonary challenge with *C. neoformans*. Control mice were left unvaccinated. Mice were euthanized at specified timepoints, and individual lungs were collected for analysis. Lung weights (A), fungal burdens (B), lung leukocytes numbers (C), CD4^+^ T cells (D) and CD8^+^ T cells (E) were determined as in Methods. Vac, vaccinated. Un/Unvac, unvaccinated. 2-Ag, vaccination with CAF01-Cda1+Cda2. 4-Ag, vaccination with CAF01-Cda1+Cda2+Blp4+Cpd1Δ. DPI, day post infection. Data are from two independent experiments, each with 3-5 mice per group. Each dot represents the value obtained from an individual mouse. CFUs are presented with individual values and medians. The dotted line in B represents the inoculum of 2 x 10^4^ yeast cells. Lung weights, and numbers of leukocytes, CD4^+^ T cells and CD8^+^ T cells are presented with means ± SEM. The results of the statistical comparison between groups are shown in S3 Table.

Analysis of *C. neoformans* CFUs in the lungs revealed that mice receiving either 2-Ag or 4-Ag vaccinations exhibited enhanced fungal control at 10 DPI, with counts approximately 8-fold lower compared to the median lung CFU of unvaccinated mice (Fig 3B). By the end of the study, all unvaccinated mice succumbed (see Fig 2A). In contrast, the majority of survivors (82.3%) from the vaccinated groups had lung CFUs below the inoculum level of 2 x 10^4^ CFU.

No significant differences in lung leukocytes were observed between unvaccinated and vaccinated groups at 0 DPI. However, augmented recruitment of lung leukocytes was evident following vaccination and infection (Fig 3C). At 10 DPI, all infected mice displayed significantly higher numbers of lung leukocytes than their corresponding groups before infection. Nonetheless, the increase in 2-Ag and 4-Ag vaccinated mice was approximately four-fold higher compared to that seen in the unvaccinated counterparts. Leukocyte numbers for vaccinated survivors at 70 DPI decreased to levels similar to those seen before infection.

Given the established importance of antigen-specific T cells for anti-cryptococcal defenses [11, 21-23], we enumerated lung CD4^+^ (Fig 3D) and CD8^+^ T cells (Fig 3E). Before infection, we did not see a difference in CD4^+^ T cell numbers between vaccinated and unvaccinated groups. At 10 DPI, CD4^+^ T cells did not remarkably increase in unvaccinated mice, but both vaccinated mouse groups had significant CD4^+^ T cell recruitment to their lungs (4-5 times higher than unvaccinated counterparts). As the infection was controlled, the CD4^+^ T cell numbers in 70-DPI survivors dramatically decreased, approaching levels comparable to those observed in naïve or vaccinated mice at 0 DPI. While a significant increase in CD8^+^ T cells was also observed following vaccination and infection, it was notably less than the increase in CD4^+^ T cells.

### Pulmonary Th1 and Th17 responses elicited by CAF01 subunit vaccination and infection

Next, we performed ex vivo experiments to further elucidate the pulmonary immune responses in the same groups of vaccinated and/or infected mice described in Fig 3. Specifically, we assessed CD154 expression as a marker of CD4^+^ T cell activation and measured antigen-stimulated intracellular production of Interferon gamma (IFNγ), tumor necrosis factor alpha (TNFα), and interleukin (IL)-17 to investigate the vaccine-induced immune response profiles of pulmonary CD4^+^ helper T cells. The antigen stimulus used was heat-killed (HK) *C. neoformans*.

In uninfected mice (0 DPI), only small numbers of lung CD4^+^ T cells expressing CD154 or producing cytokines were detected, regardless of mouse vaccination status and whether the lung cells were antigen-stimulated ex vivo (Fig 4A, 4B, 4C 4D). By 10 DPI, a substantial increase in activated CD4^+^ T cells was observed in the lungs of 2-Ag or 4-Ag vaccinated and infected mice compared to their unvaccinated counterparts (Fig 4E). However, ex vivo antigen stimulation did not further enhance CD4^+^ T cell activation in either unvaccinated or vaccinated groups. Simultaneously, lung CD4^+^ T cells from vaccinated and infected mice exhibited significantly higher production of IFNγ, TNFα, and IL-17 after HK *C. neoformans* stimulation, representing a 13∼49-fold increase compared to cells from unvaccinated infected lungs (Fig 4F, 4G, 4H). The numbers of activated and cytokine-producing CD4^+^ T cells decreased at 70 DPI for vaccinated survivors (Fig 4I, 4J, 4K, 4L) but remained substantially higher than the cell counts observed in vaccinated and uninfected mice at 0 DPI. No differences in cell activation or cytokine production were observed between the 2-Ag and 4-Ag vaccinated groups at any time points.

**Fig 4.**
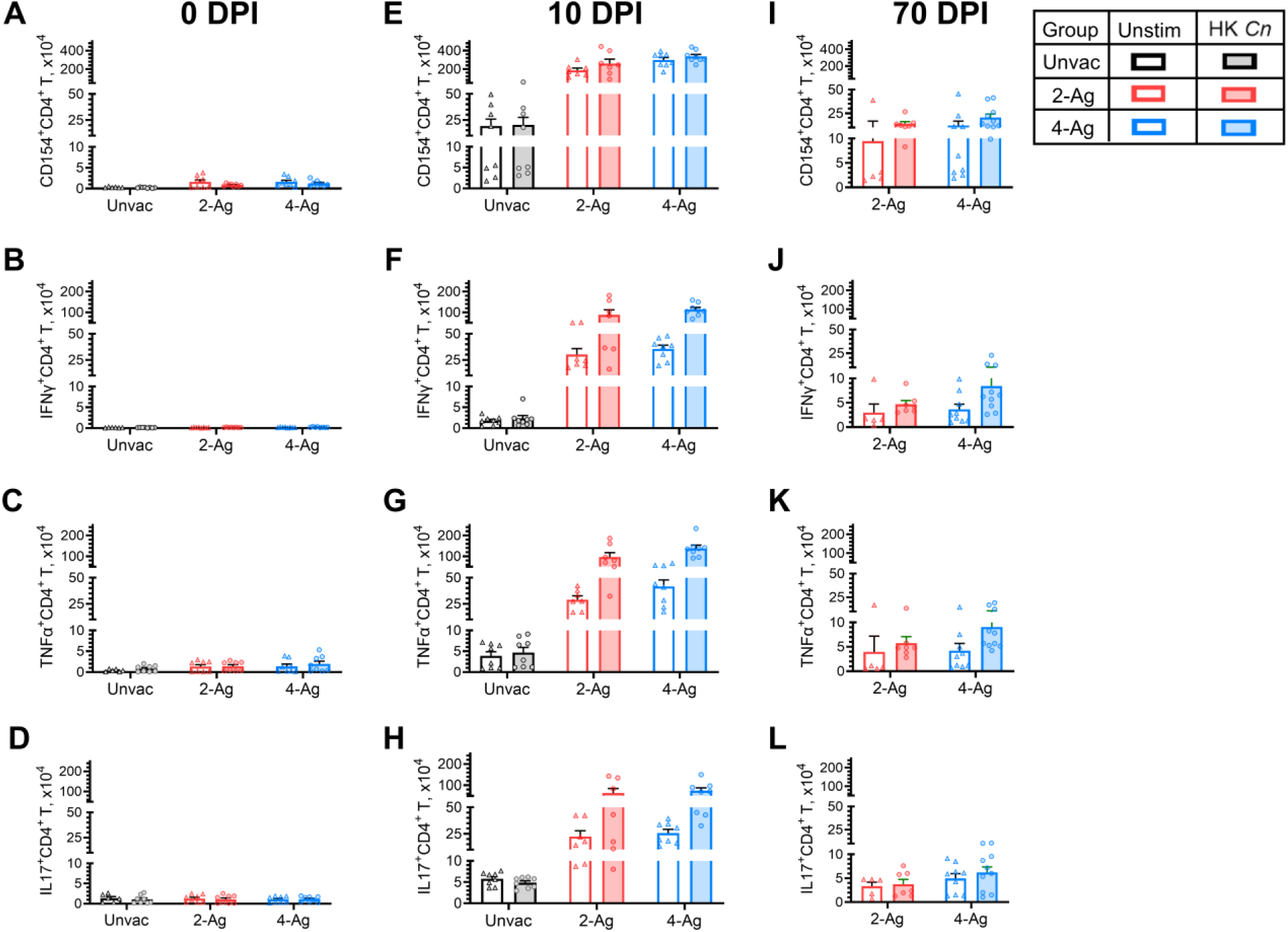
Analysis of ex vivo antigen-stimulated CD4^+^ T activation and cytokine production following CAF01-adjuvanted combination vaccination and infection. BALB/c mice were vaccinated subcutaneously thrice at biweekly intervals with either 2-Ag (Cda1+Cda2) or 4-Ag (Cda1+Cda2+Blp4+Cpd1Δ) combination vaccines adjuvanted in CAF01. Two weeks after the last boost, mice were orotracheally challenged with *C. neoformans*. Mice were euthanized at 0 DPI (uninfected), 10 DPI, or 70 DPI. Controls included unvaccinated mice euthanized at 0 DPI or 10 DPI. Lungs were harvested and single-cell suspensions were prepared. Leukocytes were cultured in complete media supplemented with amphotericin B and stimulated with HK *C. neoformans* or left unstimulated (Unstim) for 18 h. Then the cells were collected, stained, and analyzed by polychromatic FACS, as described in Methods. The numbers of CD4^+^ T cells expressing the activation marker CD154 (Fig 4A, 4E, 4I), or producing the intracellular cytokines IFNγ (Fig 4B, 4F, 4J), TNFα (Fig 4C, 4G, 4K), and IL-17 (Fig 4D, 4H, 4L) following ex vivo stimulation were determined. Unvac, unvaccinated. DPI, days post infection. HK, heat-killed. Data are from two independent experiments, each with 3-5 mice per group. Data are presented as means ± SEM. Each dot represents the value obtained from an individual mouse. Statistical comparisons between groups are shown in S4 Table.

Given the postulated importance of polyfunctional CD4^+^ T cells for vaccine-induced immunity [24], we determined the numbers of CD4^+^ T cells which produced only a single cytokine (IFNγ^+^, IL-17^+^, or TNFα^+^) versus those which produced two or three cytokines (Fig 5). These studies were performed at 10 DPI and included mice which were unvaccinated, vaccinated with 2-Ag, and vaccinated with 4-Ag. Numbers of cytokine-producing CD4^+^ T cells were low in the unvaccinated mice with double and triple positive cells infrequently seen. Populations induced by 2-Ag or 4-Ag immunization were primarily IFNγ^+^ or TNFα^+^ single-positive CD4^+^ T cells, followed by the IL-17^+^ single-positive cells and a double-positive population expressing both TNFα and IFNγ. TNFα^+^IL-17^+^ cells were also observed, albeit to a lesser extent. There were also increased numbers of IFNγ^+^IL-17^+^ double-positive and IFNγ^+^TNFα^+^IL-17^+^ triple-positive cells. Similar results were found when the data were analyzed by percentage of each subpopulation rather than actual cell numbers (see pie charts in Fig 5).

**Fig 5.**
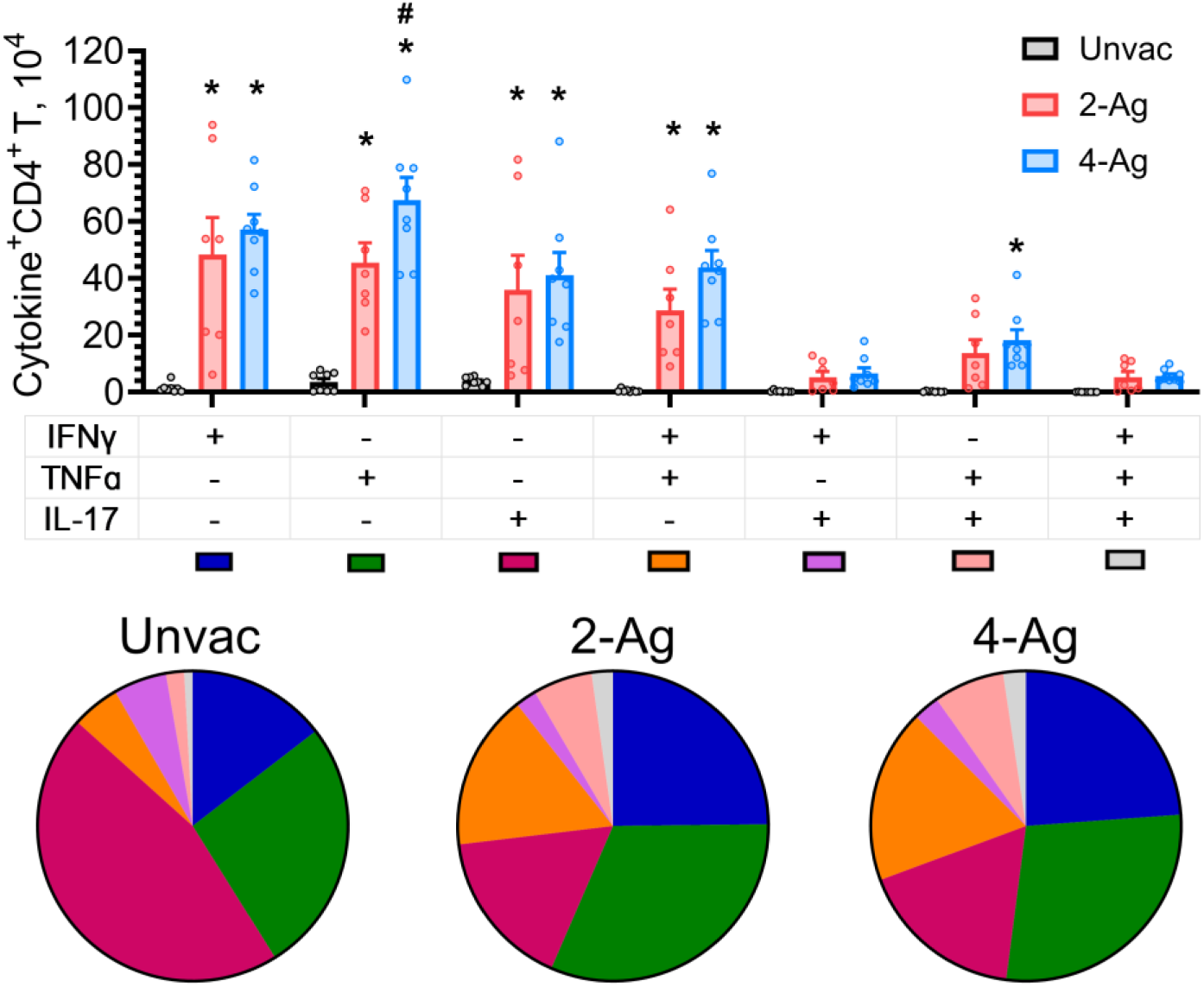
Analysis of polyfunctional lung CD4^+^ T cells following ex vivo stimulation with HK *Cryptococcus*. Lung leukocytes were prepared from the same sets of mice (10 DPI) as described in Fig 4. Leukocytes were cultured in complete media supplemented with amphotericin B and stimulated with HK *C. neoformans* for 18 h. Brefeldin A was added during the last 4 h of culture. Then the cells were collected, stained, and analyzed by polychromatic FACS. After Boolean gating analysis in FlowJo software, the frequencies of the seven possible CD4^+^ T cell subpopulations expressing any combination of the IFNγ, TNFα, and IL-17 cytokines were determined. Additionally, the corresponding numbers of each subpopulation were subsequentially enumerated. Unvac, unvaccinated. 2-Ag, vaccination with CAF01-Cda1+Cda2. 4-Ag, vaccination with CAF01-Cda1+Cda2+Blp4+Cpd1Δ. Data are from two independent experiments, each with 3-5 mice per group. Each dot represents the value obtained from an individual mouse. Data are presented as means ± SEM. The bar graphs show the cell numbers of each subpopulation. Means of cytokine-producing CD4^+^ T cells were compared. *P* < 0.05 was considered significantly different. * denotes a significant comparison with Unvac group, # denotes a significant comparison between 2-Ag and 4-Ag groups. Pie charts represent the fraction of each subpopulation in cytokine-expressing CD4^+^ T cells. Pie chart color-coding for each subpopulation is shown below the bar graph.

### Cytokine production by lung CD8^+^ T cell responses following vaccination and infection

A significant lung CD8^+^ T cell response was seen in vaccinated and infected mice, albeit not as robust of the CD4^+^ T cell response. Therefore, we examined production of IFNγ, TNFα, and IL-17 by lung CD8^+^ T cells after ex vivo stimulation with HK *C. neoformans* (Fig 6). Cytokine positive cells were infrequently observed at 0 DPI for both unvaccinated and vaccinated groups (Fig 6A, 6B, 6C). Meanwhile, we observed a sizable number of CD8^+^ T cells producing IFNγ, TNFα, IL-17 in vaccinated mice 10 DPI upon antigen stimulation (Fig 6D, 6E, 6F). CD8^+^ T cell numbers expressing IFNγ or TNFα were greater in the group vaccinated with 4-Ag compared with 2-Ag. However, the numbers of those cytokine-producing CD8^+^ T cells were considerably lower compared to their CD4^+^ T cell counterparts. At 70 DPI, vaccinated surviving mice still had increased numbers of CD8^+^ T cells producing IFNγ, but not TNFα or IL-17, following ex vivo stimulation with HK *C. neoformans* (Fig 6G, 6H, 6I).

**Fig 6.**
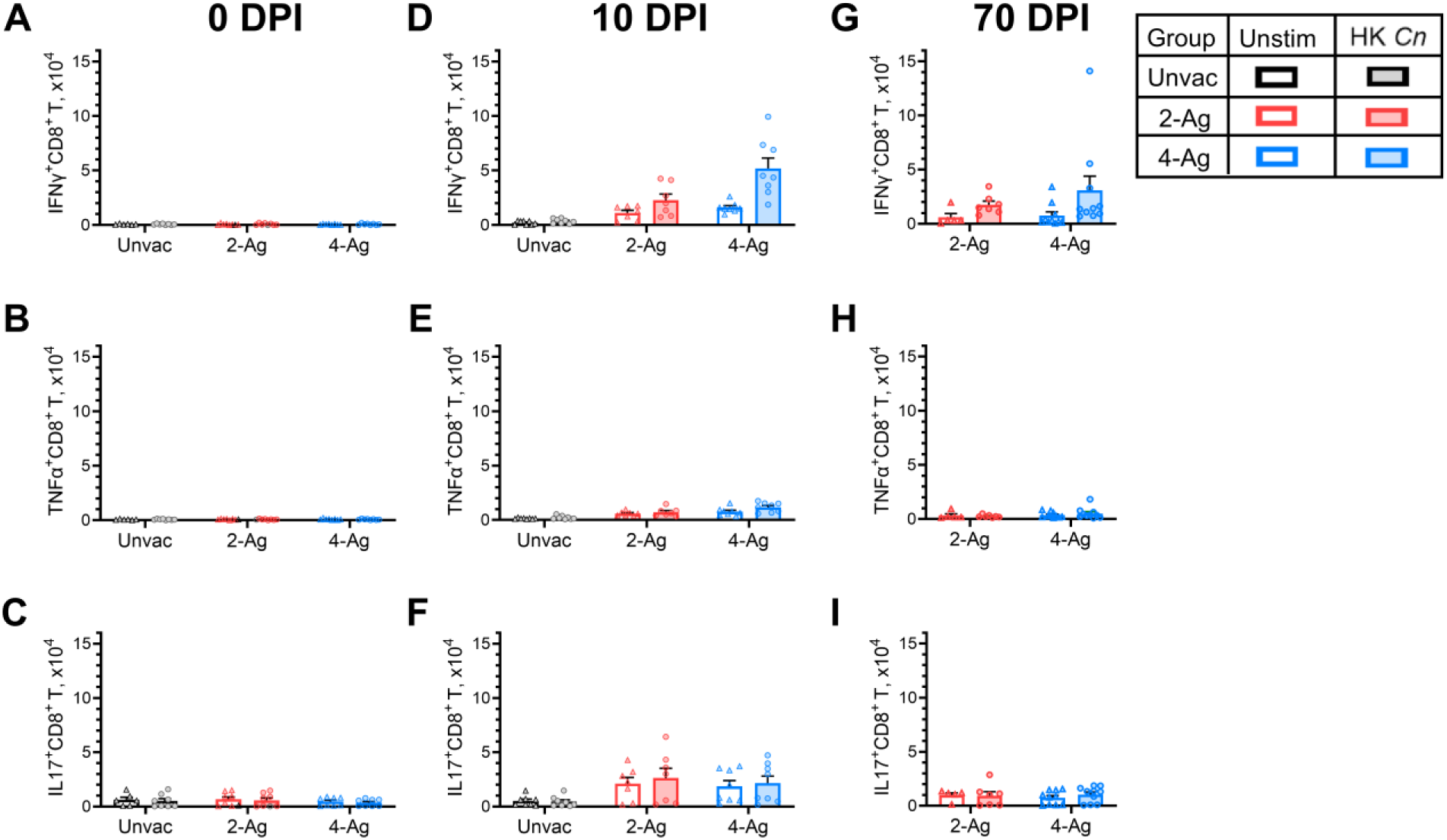
Analysis of ex vivo antigen-stimulated CD8^+^ T cytokine production following CAF01-adjuvanted combination vaccination and infection. Lung leukocytes were prepared from the same set of mice as described in Fig 4. Leukocytes were cultured in complete media supplemented with amphotericin B and stimulated with HK *C. neoformans* or left unstimulated (Unstim) for 18 h. Brefeldin A was added during the last 4 h of culture. Then the cells were collected, stained, and analyzed by polychromatic FACS. The numbers of CD8^+^ T cells producing the intracellular cytokines IFNγ (Fig 6A, 6D, 6G), TNFα (Fig 6B, 6E, 6H), and IL-17 (Fig 6C, 6F, 6I) following ex vivo stimulation were determined. Unvac, unvaccinated. DPI, days post infection. 2-Ag, vaccination with CAF01-Cda1+Cda2. 4-Ag, vaccination with CAF01-Cda1+Cda2+Blp4+Cpd1Δ. HK, heat-killed. Data are from two independent experiments, each with 3-5 mice per group, and are presented as means ± SEM. Each dot represents the value obtained from an individual mouse. Statistical comparisons between groups are shown in S5 Table.

### Antigen-specific IFNγ responses induced following vaccination and infection correlate with enhanced antifungal activity

Given the critical importance of IFNγ in host defenses against cryptococcosis [25-27], in the final set of experiments we isolated lung cells, splenocytes, and peripheral blood mononuclear cells (PBMCs) from mice which were vaccinated and/or infected, and then measured IFNγ production following ex vivo stimulation with HK *C. neoformans*. Lung leukocytes from both unvaccinated and vaccinated mice at 0 DPI showed undetectable to low levels of IFNγ production following HK *Cryptococcus* stimulation (Fig 7A). In contrast, antigen-stimulated splenocytes from 2-Ag or 4-Ag vaccinated mice exhibited a small increase in IFNγ release compared to naïve mice (Fig 7B). Meanwhile, IFNγ production by PBMCs was undetectable in all groups in the absence of infection (Fig 7C).

**Fig 7.**
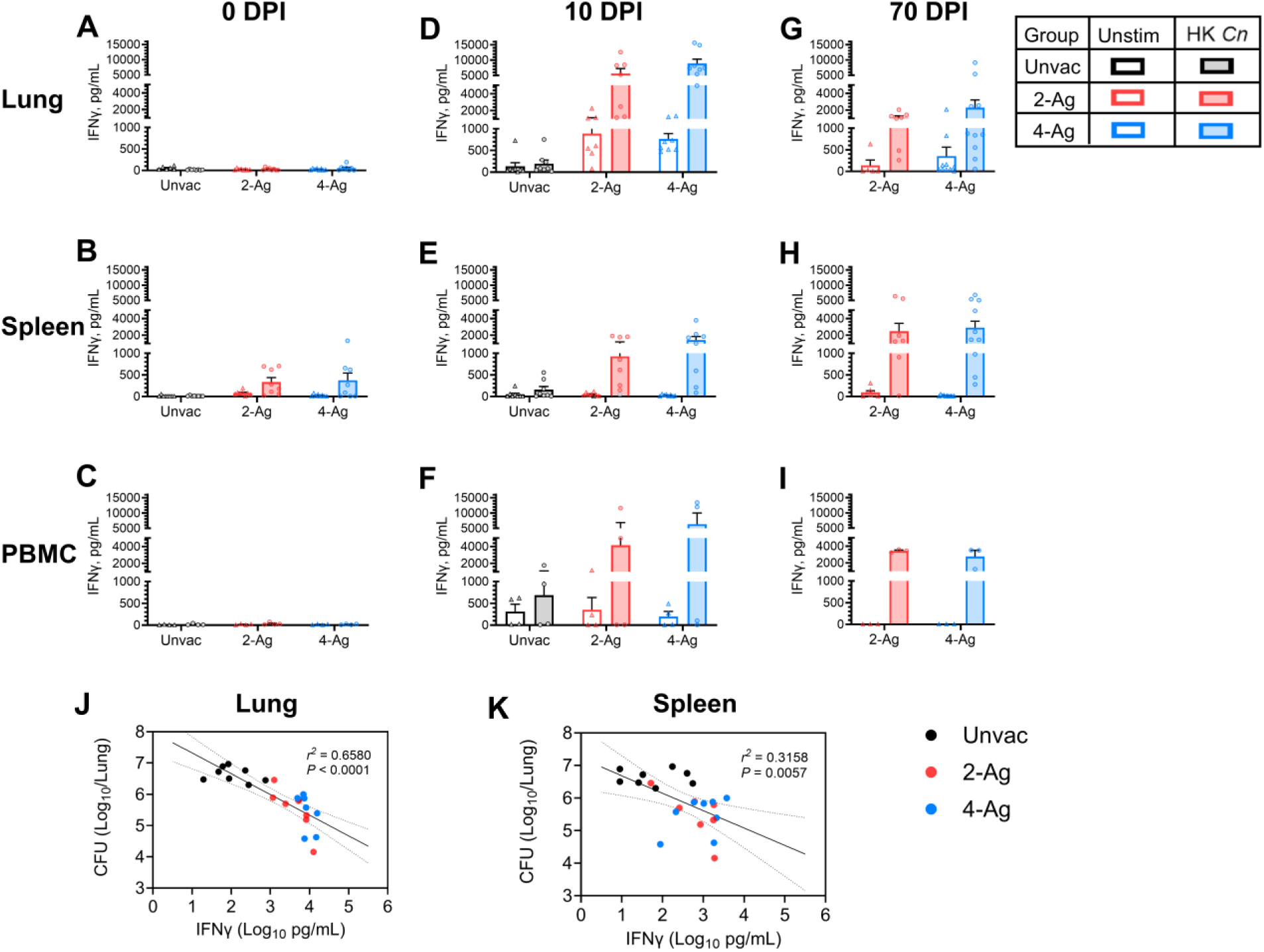
IFNγ production by ex vivo stimulated lung leukocytes, splenocytes, and PBMCs following vaccination and/or infection. BALB/c mice were vaccinated subcutaneously thrice at biweekly intervals with either 2-Ag (Cda1+Cda2) or 4-Ag (Cda1+Cda2+Blp4+Cpd1Δ) combination vaccines adjuvanted in CAF01. Two weeks after the last boost, mice were orotracheally challenged with *C. neoformans*. Mice were euthanized at 0 DPI (uninfected), 10 DPI, or 70 DPI. Controls included unvaccinated mice euthanized at 0 DPI or 10 DPI. Lung leukocytes, splenocytes and PBMCs were prepared as described in Methods. Cells were cultured in complete media supplemented with amphotericin B and stimulated with HK *C. neoformans* or left unstimulated (Unstim) for either 18 h (lung) or 3 days (spleen and PBMC). Supernatants were collected and analyzed for IFNγ by ELISA. Antigen-stimulated production for lungs (Fig 7A, 7D, 7G), spleens (Fig 7B, 7E, 7H), and PBMCs (Fig 7C, 7F, 7I) are presented as means ± SEM; statistical comparisons between groups are shown in S6 Table. Each dot represents the value obtained from an individual mouse except for PBMCs where each dot represents technical duplicates of pooled PBMCs from each independent experiment. Correlations between lung CFUs and lung leukocytes IFNγ levels (Fig 7J), or splenocytes IFNγ production (Fig 7K) following HK *Cn* stimulation were analyzed with simple linear regression (10 DPI). The results are presented with best-fit line and confidence bands. The Pearson correlation was used for statistical analysis. A *P* value < 0.05 was considered statistically significant. Unvac, unvaccinated. DPI, days post infection. HK, heat-killed. Data are from two independent experiments, each with 3-5 mice per group.

At 10 DPI, infected but not vaccinated mice had no significant antigen-specific IFNγ production in lung leukocytes (Fig 7D) or splenocytes (Fig 7E). There was, however, a modest stimulation of IFNγ in PBMCs (Fig 7F). Conversely, vaccinated mice studied at 10 DPI exhibited robust IFNγ release following HK *C. neoformans* stimulation, with the response being more pronounced for lung leukocytes (Fig 7D) and PBMCs (Fig 7F) than for splenocytes (Fig 7E).

In mice that survived until 70 DPI, recall IFNγ responses were observed following ex vivo stimulation of cells from all three anatomical sites (Fig 7G, 7H, 7I). Compared with corresponding groups at 10 DPI, IFNγ levels decreased in lung leukocytes while remaining about the same in splenocytes and PBMCs.

IFNγ levels in lung leukocytes and splenocytes following HK *Cryptococcus* stimulation inversely correlated with lung CFUs at 10 DPI (Fig 7J, 7K). Higher IFNγ production corresponded to better fungal control, with a higher correlation observed between fungal burden and lung leukocyte IFNγ response (Fig 7J) compared with splenocyte response (Fig 7K).

## Discussion

The selection of adjuvants in subunit vaccines is pivotal in contemporary vaccine development. In addition to boosting immunogenicity, adjuvants inform the nature of the immune response generated [28-30]. For example, alum, commonly used in licensed subunit vaccines, tends to promote strong antibody responses crucial for viral infection prevention but elicits relatively weak CD4^+^ T cell responses which are generally Th2-biased [31]. However, while vaccine-induced immunity requirements may differ from natural infection, Th1-biased (and to a lesser extent, Th17-biased) adaptive responses are crucial for protection against cryptococcosis [7, 32]. Thus, CAF01, which promotes Th1/Th17 responses [18, 33, 34] and has a record of safety and immunogenicity in animal and clinical studies [17-20], garnered our attention.

In our initial experiments, we explored the efficacies of four CAF01-adjuvanted monovalent subunit cryptococcal vaccines. Compared with multivalent vaccines, monovalent vaccines are easier to get clinically approved, as only one antigen needs to be manufactured and tested. We found that CAF01-adjuvanted vaccines containing either recombinant Cda1 or Cda2 provided 40-60% protection against an otherwise lethal pulmonary cryptococcal infection in both BALB/c and C57BL6 mice. Cda1 and Cda2 are chitin deacetylases responsible for deacetylating cell wall chitin to chitosan, a process which is necessary for cryptococcal virulence [35]. We previously demonstrated GP-based vaccines containing Cda1 and Cda2 significantly protected BALB/c and C57BL/6 mice [10, 11].

Blp4 is a barwin-like protein family member of unknown function whose gene is highly expressed in the cerebrospinal fluid from cryptococcal meningitis patients [36, 37]. We previously showed GP-based vaccines containing Blp4 afforded 100% and 70% survival in BALB/c and C57BL/6 mice, respectively [10]. However, the CAF01 adjuvanted formulation resulted in a nonsignificant survival extension in C57BL/6 and no survival benefit in BALB/c. The reason for the disparate results with the two adjuvants merits further study but could be due to differences in the potency of the adjuvants and the immune responses elicited. Cpd1, a carboxypeptidase family member with homology to human cathepsin A, partially protects mice from experimental cryptococcosis when delivered in GP vaccines [9, 10]. Due to concerns about molecular mimicry eliciting autoimmunity [38], we deleted the homologous region. When formulated with CAF01, the modified recombinant protein, Cpd1Δ promoted modest protection in C57BL/6 mice and had no significant efficacy in BALB/c mice. The loss of some antigen immunogenicity may be a necessary tradeoff as vaccines containing proteins with human homologs generally should be avoided [38].

Taken together, our data revealed that CAF01, when combined with selectively chosen single antigens, afforded partial protection against cryptococcal infection. To improve the efficacy, we next tested CAF01 vaccines formulated with multiple antigens. With other vaccines, increasing the number of antigens expands the repertoire of antigen-specific immune responses [39-41]. This is particularly pertinent for human T cell vaccines given the diversity of human leukocyte antigen (HLA) class II alleles in the population [42]. Moreover, multivalent vaccines mitigate the risk of immune evasion by pathogens by reducing the likelihood of the emergence of escape mutants [43]. In our study, the CAF01-adjuvanted bivalent or quadrivalent vaccines significantly enhanced the protection against lethal pulmonary infection of *C. neoformans* in both BALB/c and C57BL/6 mice. Notably, mice vaccinated with the 4-Ag vaccines were 100% protected until the end of the study. Weight loss was observed in mice that received the multivalent vaccines during the early phase of infection, which we speculate is due to early onset of an inflammatory, but beneficial immune response against the invading pathogen. Notably though, the vaccinated mice regained the lost weight while the unvaccinated mice had late onset weight loss followed by death.

Interestingly, the protection elicited by the multivalent vaccines was completely abolished in CARD9^-/-^ mice. This finding is consistent with the role of CAF01 as a Mincle agonist. Upon ligand binding, Mincle can activate the CARD-9 downstream signaling pathway, leading to the production of pro-inflammatory cytokines and the initiation of antifungal immune responses [13]. However, the involvement of other pathogen recognition receptors cannot be ruled out in our study, as Dectin-1, Dectin-2, and some toll-like receptors also signal through CARD9 pathways [44]. Testing vaccine efficacy in mice deficient in Mincle expression could provide a clearer understanding of the contribution of Mincle to the efficacy of our CAF01-adjuvanted vaccines.

Numerous studies have underscored the pivotal role of CD4^+^ T cells in the immune response against cryptococcal infections, highlighting their significance in both controlling the infection and in vaccine-mediated protection. The loss of CD4^+^ T cells, as notably occurs in persons with HIV/AIDS, markedly increases susceptibility to cryptococcosis [1]. CD4^+^ T cells, particularly those differentiating into Th1 and Th17 phenotypes, are critical orchestrators of effective immune defenses against *Cryptococcus* [11, 45, 46]. Additionally, vaccine-induced protection is lost in mice with a genetic deficiency in CD4^+^ T cells or following CD4^+^ T cell depletion [11, 47]. In the current study, compared with unvaccinated mice, lungs from mice vaccinated with CAF01 multivalent vaccines had fewer CFUs yet weighed more and had greater numbers of leukocytes when analyzed 10 DPI. The leukocyte influx featured a robust recruitment of activated CD4^+^ T cells to the lungs. Moreover, by multicolor flow cytometry, many of the CD4^+^ T cells harvested from vaccinated and infected lungs produced IFNγ, TNFα (Th1), and IL-17 (Th17) following antigenic stimulation ex vivo. Many of the CD4^+^ T cells produced multiple cytokines which may be important as the presence of polyfunctional antigen-specific CD4^+^ T cells correlates with immune protection [24, 48]. Collectively, our data demonstrate that CAF01-adjuvanted multivalent subunit vaccines induce robust pulmonary Th1 and Th17 immune responses, leading to efficient control of *C. neoformans* in mice.

In comparison, although there was a significant increase in CD8^+^ T cells in vaccinated and infected mice, their numbers were considerably lower compared with those of infiltrating CD4^+^ T cells. This observation with CAF01 subunit vaccines is consistent with our previous findings that CD4^+^ responses, but not CD8^+^ T cell responses, play a non-redundant role after GP vaccination and cryptococcal infection [11]. Notably, in the CAF01-4Ag vaccine group, the mice had much higher numbers of cytotoxic CD8^+^ T (Tc1) cells than in unvaccinated or 2-Ag vaccinated mice.

As Tc1 cells have a direct killing effect on *Cryptococcus* [49], this could additionally contribute to the complete protection observed in mice with CAF01-4Ag vaccination. Moreover, in situations where CD4^+^ T cell function is impaired, such as in advanced HIV infection, CD8^+^ T cells might play a more dominant role.

In addition to Th1 cells, IFNγ is made by other cell types including natural killer (NK) cells, dendritic (DC) cells and Tc1 cells [50]. IFNγ activates macrophages and induces production of nitric oxide and reactive oxygen species to facilitate the efficient killing of *Cryptococcus* [51]. GP vaccine-mediated protection against cryptococcosis was lost in IFNγ and IFNγ receptor knock-out mice [11]. We investigated IFNγ secretion following ex vivo antigenic stimulation of lung leukocytes, splenocytes, and PBMCs obtained from vaccinated and/or infected mice. We observed the production of IFNγ from splenocytes, but not lung leukocytes or PBMCs in mice which were vaccinated but not infected. In contrast, IFNγ responses were prominent in all three compartments after vaccination and infection, and were durable until 10 weeks post-infection when the mice had largely cleared the infection. Moreover, IFNγ levels, especially in the lung compartment, were positively correlated with fungal clearance. Thus, our data add evidence to the indispensable role of IFNγ during cryptococcal infection and vaccination.

In summary, our preclinical studies provide a rationale for the use of CAF01-adjuvanted subunit *Cryptococcus* vaccines. While monovalent vaccination with cryptococcal recombinant proteins partially protected different mouse strains, the protection was greatly augmented when vaccination with multivalent antigens was adopted. Given population variations in HLA alleles and heterogeneity among clinical cryptococcal isolates, we believe effective subunit cryptococcal vaccines will need to be multivalent. Importantly, CAF01 subunit vaccination also elicited robust Th1 and Th17 immune responses following infection, and IFNγ responses correlated with fungal control as measured by lung CFUs. To pave the way for clinical trials, future studies in our laboratory will optimize antigen combinations in the CAF01 vaccine formulations, optimize dose and vaccine schedules, and test efficacy against genetically diverse clinical *Cryptococcus* isolates.

## Materials and Methods

### Sex as a biological variable

#### Ethics statement

All experimental procedures were conducted in accordance with guidelines approved by the Institutional Animal Care and Use Committee of UMCMS (#202100034).

### Mice

Six- to 10-week-old BALB/c (JAX stock #000651), C57BL/6 (JAX stock #000664) mice, and the CARD9 knock-out mice on the C57BL/6 background (CARD9^-/-^, JAX stock #028652)[52] were purchased from the Jackson Laboratory (Bar Harbor, ME). Our study examined both male and female mice in approximately equal numbers, and similar findings are reported for both sexes.

Mice were housed and maintained at the animal facilities of University of Massachusetts Chan Medical School (UMCMS) under specific-pathogen-free conditions. House bred mice were also used for some of the experiments.

### CAF01 adjuvanted vaccines and immunization schedules

The cryptococcal antigens Cda1, Cda2, Blp4, and Cpd1Δ were expressed as His-tagged recombinant proteins in *E. coli* and purified as described [9, 11]. Purified protein (7 - 10 mg/mL) in 6M Urea, 20 mM Tris, pH 7.9 was stored at -80°C. More detailed information regarding these four recombinant proteins is shown in S1 Table. As noted above, Cda1, Cda2, Blp4, and Cpd1 confer protection in both BALB/c and C57BL/6 when adjuvanted with GPs [10]. Cpd1Δ is a version of Cpd1 deleted of amino acids from the region with homology to human cathepsin.

The liposomal adjuvant CAF01, composed of 2500 µg/mL DDA and 500 µg/mL TDB, as well as Tris Buffer (10 mM, pH 7.0, supplemented with 2% glycerol), was supplied by Statens Serum Institut (Copenhagen, Denmark). To formulate a dose of vaccine, 1 µL of antigen was diluted with Tris Buffer to a final volume 50 µL and briefly vortexed. The antigen-Tris Buffer solution was then combined with 50 µL of CAF01 and vortexed. Thus, each injection had a final volume of 100 µL. Three types of vaccine formulations were devised: (i) A single antigen vaccine, consisting of one injection with 7 - 10 µg of cryptococcal protein. (ii) A CAF01-Cda1/Cda2 combination vaccine, comprised of two injections, with each injection containing 10 µg of either Cda1 or Cda2. (iii) A 4-antigen combination vaccination (CAF01-Cda1/Cda2/Blp4/Cpd1Δ) consisting of two injections; one injection contained 5 µg each of Cda1 and Cda2, while the other contained 5 µg each of Blp4 and Cpd1Δ.

Mice were immunized via the subcutaneous route, either at the midline of the abdomen for single injection or both sides, about 1 cm from the midline for double injections. The primary immunization was followed by two booster doses (formulated and administered the same way) at two-week intervals between each vaccination. As controls, some mice were left unvaccinated, while others received CAF01 without antigens (three 100 µL doses that contained 50 µL of CAF01 and 50 µL of Tris Buffer).

### *C. neoformans* culture and infection

Two weeks following the final vaccination, mice were subjected to orotracheal challenge using the highly virulent *C. neoformans* strain KN99ɑ (hereafter referred to as KN99), as in our prior studies[11, 53]. Briefly, KN99 recovered from a glycerol stock stored at -80°C was grown on yeast-peptone-dextrose (YPD) agar for 48 hours at 30°C and then cultured overnight in liquid YPD at 30°C. The yeast cells were harvested, washed, counted, and resuspended in phosphate buffered saline (PBS). Each mouse received a 50 µL inoculum under anesthesia, with BALB/c mice receiving 2 x 10^4^ yeast cells and C57BL/6 and CARD9^-/-^ mice receiving 1 x 10^4^ yeast cells. CFUs of the inoculum were enumerated by culturing on Sabouraud dextrose agar (Thermo Fisher, Pittsburgh, PA).

Following the challenge, mice were monitored daily until death. Mice that reached humane endpoints and mice that survived 70 DPI were euthanized with CO_2_ asphyxiation. In specific groups, body weight measurements were recorded at least biweekly, and mice were euthanized at indicated timepoints (0 DPI, 10 DPI, or 70 DPI), to collect organs for subsequent ex vivo experiments.

### Sample collection, cell preparation and lung CFU enumeration

After inducing anesthesia with isoflurane, cardiac puncture was performed with heparinized syringes to collect blood, which was subsequently pooled within the respective experimental groups. Following dilution with an equal volume of PBS containing 2% fetal bovine serum (FBS, Thermo Fisher), the pooled blood was carefully layered over 5 mL of Ficoll-Paque PREMIUM 1.084 (Cytiva, Uppsala, Sweden) in 15 mL SepMate-50 tubes (STEM Cell Technology, Cambridge, MA). PBMCs were collected from the interphase layer after centrifugation at 1200xg for 30 min.

Spleens and lungs were collected and processed individually after the removal of blood. Spleens were gently pressed against a 70 µm cell strainer using the plunger of a 3 mL syringe. Single cells were obtained by flushing the cell strainer with 6 mL complete medium (RPMI 1640 supplemented 10% FBS, 1% HEPES, 1% GlutaMAX, and 1% Penicillin–Streptomycin, all purchased from ThermoFisher Scientific).

Lungs were rinsed with PBS. Lung cells were prepared using the MACS Lung Dissociation Kit for mice, following the manufacturer’s instructions (Miltenyi Biotec, Bergisch Gladbach, Germany). A 200 µL portion of each lung digest was designated for lung fungal burden assessment, with both undiluted and diluted samples cultured on Sabouraud dextrose agar. Following a 2-day incubation at 30°C, CFUs were enumerated, with a lower limit of detection of 20 CFU/lung. The remaining lung samples were passed through a 70 µm cell strainer. Leukocytes from lung single-cell suspensions were enriched using a Percoll gradient (67% and 40%, Cytiva; centrifuged at 800xg for 20 minutes without brake), followed by the collection of cells from the interphase. Single cells obtained from PBMCs, splenocytes and lung leukocytes were washed and resuspended in complete medium. Cell concentration and viability were then determined with Trypan Blue stain and a T20 cell counter (Bio-Rad, Hercules, CA).

### Ex Vivo Culture and Stimulation

PBMCs (2 x 10^5^ cells/well), splenocytes (1 x 10^6^ cells/well), and lung leukocytes (4 x 10^5^ cells/well) were cultured in round-bottomed 96-well plates, each containing 200 µL of complete culture medium supplemented with 0.5 µg/mL amphotericin B (Thermo Fisher). The cells were left unstimulated or stimulated with HK *C. neoformans* strain KN99 (50 µg/mL, dry weight, prepared according to established protocols [11]). The cultures were maintained at 37°C within a controlled humidified atmosphere supplemented with 5% CO_2_.

Following a 3-day incubation, supernatants from cultured PBMCs and splenocytes were harvested for determination of IFNγ concentration. Meanwhile, the lung leukocytes were cultured for 18 hours, with the addition of brefeldin A (5 µg/mL, BioLegend, San Diego, CA) during the last 4 hours of incubation. Supernatants were subsequently collected for IFNγ quantification, and the lung leukocytes were harvested for staining, followed by analysis using flow cytometry (FACS). Each sample was prepared and analyzed in duplicate.

### Intracellular Staining for Cytokines

Harvested lunCg leukocytes were washed with PBS, followed by viability staining with LIVE/DEAD green fixable dead cell stain kit (1:1000 dilution, ThermoFisher). For surface staining, cells were incubated with anti-mouse CD3-PE, CD4-PerCP/Cyanine5.5 and CD8-APC antibodies in the dark at 4°C for 30 min in 50 µL FACS buffer (PBS with 0.5% bovine serum albumin). Cells were then fixed and permeabilized with the Intracellular Fixation & Permeabilization Buffer Set (ThermoFisher). Rat anti-mouse CD16/CD32 monoclonal antibody was co-incubated with cells to block Fc receptors, before intracellular staining with CD154-PE/Cyanine7, IFNγ-BV650, TNFα-APC/Cyanine7 and IL-17A-BV510 for 30min. Stained cells were resuspended in FACS buffer and acquired with a 5-laser Bio-Rad ZE5 flow cytometer (Bio-Rad). FACS data were analyzed using FlowJo version 10.9 software (BD Biosciences, Franklin Lakes, NJ). Antibodies used for staining are listed in S2 Table. Gating strategies, as demonstrated in S1 Fig, utilized fluorescence minus one controls and isotype controls.

### Quantification of IFNγ production

All culture supernatants were promptly frozen and stored at -80°C until analyzed. Samples were thawed, diluted 1:2, and IFNγ levels were measured with R&D Systems Mouse IFNγ DuoSet ELISA Kit (Bio-Techne, Minneapolis, MN) in accordance with the manufacturer’s instructions. The detection limit was 10 pg/mL; samples below the detection limit were assigned a value of 9 pg/mL.

### Statistics

GraphPad Prism, version 10.1.2 (GraphPad Software, La Jolla, CA) was used for data analysis and graphical representation. In survival studies, Kaplan-Meier survival curves underwent statistical assessment for significance utilizing the Mantel-Cox log-rank test. Median survival differences ≤ 3 days were deemed not biologically meaningful and, consequently, were not assessed for statistical significance. Comparison of lung CFUs was executed through the nonparametric Mann-Whitney test as the data did not follow a Poisson distribution. Meanwhile, evaluation of normally distributed data such as lung weights, lung leukocytes, CD4^+^ T cell, and CD8^+^ T cell numbers among groups employed the one-way ANOVA test with the Šídák correction for multiple comparisons.

For ex vivo intracellular staining experiments, lungs were analyzed individually, and the data presented are the averages of technical duplicates for each sample. Cytokine expression comparisons across groups were performed using the two-way ANOVA test with Tukey or Šídák correction applied for multiple comparisons. For IFNγ ELISA experiments, data were analyzed for the lungs and spleens of individual mice, with averages of technical duplicates presented. Meanwhile, technical duplicates of pooled PBMCs for each group were presented and analyzed. The two-way ANOVA test with Tukey or Šídák correction was applied for multiple comparisons. Significance was defined as a *P*-value < 0.05 after corrections for multiple comparisons.

## Data Availability

All relevant data supporting the findings of this study are available within the article and supporting information..

## Author contributions

RW, LVNO, DCh, CAS. and SML conceived and designed experiments. RW, LVNO, DCa, MMH and CAS performed experiments. RW, LVNO, and CAS analyzed the data. RW, CAS, and SML wrote the paper. CAS and SML are co-corresponding authors for the article. All authors contributed to the manuscript and gave their approval for the final version to be submitted.

## Acknowledgements

The authors extend their gratitude to Dr. Ashraf Ibrahim for recommending the evaluation of CAF01 in cryptococcal vaccine studies and to Dr. Gabriel Kristian Pedersen (Statens Serum Institut) for assistance providing the CAF01. The authors thank Christina Gomez and Zhongming Mou for their help with the mouse studies.

## References

1. Williamson PR, Jarvis JN, Panackal AA, Fisher MC, Molloy SF, Loyse A, et al. Cryptococcal meningitis: epidemiology, immunology, diagnosis and therapy. Nat Rev Neurol. 2017;13(1):13–24. doi: 10.1038/nrneurol.2016.167.

2. Ngan NTT, Flower B, Day JN. Treatment of cryptococcal meningitis: how have we got here and where are we going? Drugs. 2022;82(12):1237–49. doi: 10.1007/s40265-022-01757-5.

3. World Health Organization. Guidelines for diagnosing, preventing and managing cryptococcal disease among adults, adolescents and children living with HIV. Updated June 17, 2022. Available from: https://apps.who.int/iris/bitstream/handle/10665/357088/9789240052178-eng.pdf?sequence=1..

4. Rajasingham R, Govender NP, Jordan A, Loyse A, Shroufi A, Denning DW, et al. The global burden of HIV-associated cryptococcal infection in adults in 2020: a modelling analysis. Lancet Infect Dis. 2022;22(12):1748–55. doi: 10.1016/S1473-3099(22)00499-6.

5. Loyse A, Burry J, Cohn J, Ford N, Chiller T, Ribeiro I, et al. Leave no one behind: Response to new evidence and guidelines for the management of cryptococcal meningitis in low-income and middle-income countries. Lancet Infect Dis. 2019;19(4):e143–e7. doi: 10.1016/S1473-3099(18)30493-6.

6. World Health Organization. WHO fungal priority pathogens list to guide research, development and public health action. Updated October 25, 2022. Available from: https://www.who.int/publications/i/item/9789240060241.

7. Oliveira LVN, Wang R, Specht CA, Levitz SM. Vaccines for human fungal diseases: close but still a long way to go. NPJ Vaccines. 2021;6(1):33. doi: 10.1038/s41541-021-00294-8.

8. Abraham A, Ostroff G, Levitz SM, Oyston PCF. A novel vaccine platform using glucan particles for induction of protective responses against Francisella tularensis and other pathogens. Clin Exp Immunol. 2019;198(2):143–52. doi: 10.1111/cei.13356.

9. Hester MM, Oliveira LVN, Wang R, Mou Z, Lourenco D, Ostroff GR, et al. Cross-reactivity between vaccine antigens from the chitin deacetylase protein family improves survival in a mouse model of cryptococcosis. Front Immunol. 2022;13:1015586. doi: 10.3389/fimmu.2022.1015586.

10. Hester MM, Lee CK, Abraham A, Khoshkenar P, Ostroff GR, Levitz SM, et al. Protection of mice against experimental cryptococcosis using glucan particle-based vaccines containing novel recombinant antigens. Vaccine. 2020;38(3):620–6. doi: 10.1016/j.vaccine.2019.10.051.

11. Wang R, Oliveira LVN, Lourenco D, Gomez CL, Lee CK, Hester MM, et al. Immunological correlates of protection following vaccination with glucan particles containing Cryptococcus neoformans chitin deacetylases. npj Vaccines. 2023;8(1). doi: 10.1038/s41541-023-00606-0.

12. Christensen D. Development and evaluation of CAF01. In: Schijns VEJC, O’Hagan DT, editors. Immunopotentiators in Modern Vaccines (Second Edition): Academic Press; 2017. p. 333-45.

13. Pedersen GK, Andersen P, Christensen D. Immunocorrelates of CAF family adjuvants. Semin Immunol. 2018;39:4–13. doi: 10.1016/j.smim.2018.10.003.

14. Tsoni SV, Brown GD. beta-Glucans and dectin-1. Ann N Y Acad Sci. 2008;1143:45–60. doi: 10.1196/annals.1443.019.

15. Williams SJ. Sensing lipids with mincle: structure and function. Front Immunol. 2017;8:1662. doi: 10.3389/fimmu.2017.01662.

16. Knudsen NP, Olsen A, Buonsanti C, Follmann F, Zhang Y, Coler RN, et al. Different human vaccine adjuvants promote distinct antigen-independent immunological signatures tailored to different pathogens. Sci Rep. 2016;6(1):19570. doi: 10.1038/srep19570.

17. Abraham S, Juel HB, Bang P, Cheeseman HM, Dohn RB, Cole T, et al. Safety and immunogenicity of the chlamydia vaccine candidate CTH522 adjuvanted with CAF01 liposomes or aluminium hydroxide: a first-in-human, randomised, double-blind, placebo-controlled, phase 1 trial. Lancet Infect Dis. 2019;19(10):1091–100. doi: 10.1016/S1473-3099(19)30279-8.

18. van Dissel JT, Joosten SA, Hoff ST, Soonawala D, Prins C, Hokey DA, et al. A novel liposomal adjuvant system, CAF01, promotes long-lived Mycobacterium tuberculosis-specific T-cell responses in human. Vaccine. 2014;32(52):7098-107. doi: 10.1016/j.vaccine.2014.10.036.

19. Fomsgaard A, Karlsson I, Gram G, Schou C, Tang S, Bang P, et al. Development and preclinical safety evaluation of a new therapeutic HIV-1 vaccine based on 18 T-cell minimal epitope peptides applying a novel cationic adjuvant CAF01. Vaccine. 2011;29(40):7067–74. doi: 10.1016/j.vaccine.2011.07.025.

20. Dejon-Agobe JC, Ateba-Ngoa U, Lalremruata A, Homoet A, Engelhorn J, Nouatin OP, et al. Controlled human malaria infection of healthy adults with lifelong malaria exposure to assess safety, immunogenicity, and efficacy of the asexual blood stage malaria vaccine candidate GMZ2. Clin Infect Dis. 2019;69(8):1377–84. doi: 10.1093/cid/ciy1087.

21. Chen J, Shao J, Dai M, Fang W, Yang YL. Adaptive immunology of Cryptococcus neoformans infections-an update. Front Immunol. 2023;14:1174967. doi: 10.3389/fimmu.2023.1174967.

22. Uicker WC, McCracken JP, Buchanan KL. Role of CD4+ T cells in a protective immune response against Cryptococcus neoformans in the central nervous system. Med Mycol. 2006;44(1):1–11. doi.

23. Lindell DM, Moore TA, McDonald RA, Toews GB, Huffnagle GB. Generation of antifungal effector CD8+ T cells in the absence of CD4+ T cells during Cryptococcus neoformans infection. J Immunol. 2005;174(12):7920–8. doi: 10.4049/jimmunol.174.12.7920.

24. Derrick SC, Yabe IM, Yang A, Morris SL. Vaccine-induced anti-tuberculosis protective immunity in mice correlates with the magnitude and quality of multifunctional CD4 T cells. Vaccine. 2011;29(16):2902–9. doi: 10.1016/j.vaccine.2011.02.010.

25. Wormley FL, Jr., Perfect JR, Steele C, Cox GM. Protection against cryptococcosis by using a murine gamma interferon-producing Cryptococcus neoformans strain. Infect Immun. 2007;75(3):1453–62. doi: 10.1128/iai.00274-06.

26. Leopold Wager CM, Hole CR, Campuzano A, Castro-Lopez N, Cai H, Caballero Van Dyke MC, et al. IFN-gamma immune priming of macrophages in vivo induces prolonged STAT1 binding and protection against Cryptococcus neoformans. PLoS Pathog. 2018;14(10):e1007358. doi: 10.1371/journal.ppat.1007358.

27. Pappas PG, Bustamante B, Ticona E, Hamill RJ, Johnson PC, Reboli A, et al. Recombinant interferon-gamma 1b as adjunctive therapy for AIDS-related acute cryptococcal meningitis. J Infect Dis. 2004;189(12):2185–91. doi: 10.1086/420829.

28. Pulendran B, S.Arunachalam P, O’Hagan DT. Emerging concepts in the science of vaccine adjuvants. Nat Rev Drug Discov. 2021;20(6):454–75. doi: 10.1038/s41573-021-00163-y.

29. Zhao T, Cai Y, Jiang Y, He X, Wei Y, Yu Y, et al. Vaccine adjuvants: mechanisms and platforms. Signal Transduct Target Ther. 2023;8(1):283. doi: 10.1038/s41392-023-01557-7.

30. Levitz SM, Golenbock DT. Beyond empiricism: informing vaccine development through innate immunity research. Cell. 2012;148(6):1284–92. doi: 10.1016/j.cell.2012.02.012.

31. Fan J, Jin S, Gilmartin L, Toth I, Hussein WM, Stephenson RJ. Advances in infectious disease vaccine adjuvants. Vaccines (Basel). 2022;10(7). doi: 10.3390/vaccines10071120.

32. Elsegeiny W, Marr KA, Williamson PR. Immunology of cryptococcal infections: Developing a rational approach to patient therapy. Front Immunol. 2018;9:651. doi: 10.3389/fimmu.2018.00651.

33. Lindenstrom T, Woodworth J, Dietrich J, Aagaard C, Andersen P, Agger EM. Vaccine-induced Th17 cells are maintained long-term postvaccination as a distinct and phenotypically stable memory subset. Infect Immun. 2012;80(10):3533–44. doi: 10.1128/IAI.00550-12.

34. Lindenstrom T, Agger EM, Korsholm KS, Darrah PA, Aagaard C, Seder RA, et al. Tuberculosis subunit vaccination provides long-term protective immunity characterized by multifunctional CD4 memory T cells. J Immunol. 2009;182(12):8047–55. doi: 10.4049/jimmunol.0801592.

35. Upadhya R, Lam WC, Hole CR, Parchment D, Lee CK, Specht CA, et al. Cryptococcus neoformans Cda1 and Cda2 coordinate deacetylation of chitin during infection to control fungal virulence. Cell Surf. 2021;7:100066. doi: 10.1016/j.tcsw.2021.100066.

36. Chen Y, Toffaletti DL, Tenor JL, Litvintseva AP, Fang C, Mitchell TG, et al. The Cryptococcus neoformans transcriptome at the site of human meningitis. mBio. 2014;5(1):e01087–13. doi: 10.1128/mBio.01087-13.

37. Yu CH, Sephton-Clark P, Tenor JL, Toffaletti DL, Giamberardino C, Haverkamp M, et al. Gene expression of diverse Cryptococcus isolates during infection of the human central nervous system. mBio. 2021;12(6):e0231321. doi: 10.1128/mBio.02313-21.

38. Segal Y, Shoenfeld Y. Vaccine-induced autoimmunity: the role of molecular mimicry and immune crossreaction. Cellular & Molecular Immunology. 2018;15(6):586–94. doi: 10.1038/cmi.2017.151.

39. Scott NR, Mann B, Tuomanen EI, Orihuela CJ. Multi-valent protein hybrid pneumococcal vaccines: A strategy for the next generation of vaccines. Vaccines (Basel). 2021;9(3). doi: 10.3390/vaccines9030209.

40. Uno N, Ross TM. Multivalent next generation influenza virus vaccines protect against seasonal and pre-pandemic viruses. Sci Rep. 2024;14(1):1440. doi: 10.1038/s41598-023-51024-0.

41. Chen A, Mann B, Gao G, Heath R, King J, Maissoneuve J, et al. Multivalent pneumococcal protein vaccines comprising pneumolysoid with epitopes/fragments of CbpA and/or PspA elicit strong and broad protection. Clin Vaccine Immunol. 2015;22(10):1079–89. doi: 10.1128/CVI.00293-15.

42. Stern LJ, Calvo-Calle JM. HLA-DR: molecular insights and vaccine design. Curr Pharm Des. 2009;15(28):3249–61. doi: 10.2174/138161209789105171.

43. Korber B, Hraber P, Wagh K, Hahn BH. Polyvalent vaccine approaches to combat HIV-1 diversity. Immunol Rev. 2017;275(1):230–44. doi: 10.1111/imr.12516.

44. Liu X, Jiang B, Hao H, Liu Z. CARD9 signaling, inflammation, and diseases. Front Immunol. 2022;13:880879. doi: 10.3389/fimmu.2022.880879.

45. Jarvis JN, Casazza JP, Stone HH, Meintjes G, Lawn SD, Levitz SM, et al. The phenotype of the Cryptococcus-specific CD4+ memory T-cell response is associated with disease severity and outcome in HIV-associated cryptococcal meningitis. J Infect Dis. 2013;207(12):1817–28. doi: 10.1093/infdis/jit099.

46. Wang Y, Wang K, Masso-Silva JA, Rivera A, Xue C. A heat-killed Cryptococcus mutant strain induces host protection against multiple invasive mycoses in a murine vaccine model. MBio. 2019;10(6). doi: 10.1128/mBio.02145-19.

47. Pham T, Li Y, Watford W, Lin X. Vaccination with a ZNF2(oe) strain of Cryptococcus provides long-lasting protection against cryptococcosis and is effective in immunocompromised hosts. Infect Immun. 2023:e0019823. doi: 10.1128/iai.00198-23.

48. Boyd A, Almeida JR, Darrah PA, Sauce D, Seder RA, Appay V, et al. Pathogen-specific T Cell polyfunctionality is a correlate of T Cell efficacy and immune protection. PLoS One. 2015;10(6):e0128714. doi: 10.1371/journal.pone.0128714.

49. Ma LL, Spurrell JC, Wang JF, Neely GG, Epelman S, Krensky AM, et al. CD8 T cell- mediated killing of Cryptococcus neoformans requires granulysin and is dependent on CD4 T cells and IL-15. J Immunol. 2002;169(10):5787–95. doi: 10.4049/jimmunol.169.10.5787.

50. Kak G, Raza M, Tiwari BK. Interferon-gamma (IFN-gamma): Exploring its implications in infectious diseases. Biomol Concepts. 2018;9(1):64–79. doi: 10.1515/bmc-2018-0007.

51. Leopold Wager CM, Hole CR, Wozniak KL, Wormley FL, Jr. Cryptococcus and phagocytes: complex interactions that influence disease outcome. Front Microbiol. 2016;7:105. doi: 10.3389/fmicb.2016.00105.

52. Hsu YM, Zhang Y, You Y, Wang D, Li H, Duramad O, et al. The adaptor protein CARD9 is required for innate immune responses to intracellular pathogens. Nat Immunol. 2007;8(2):198–205. doi: 10.1038/ni1426.

53. Specht CA, Lee CK, Huang H, Tipper DJ, Shen ZT, Lodge JK, et al. Protection against Experimental Cryptococcosis following Vaccination with Glucan Particles Containing Cryptococcus Alkaline Extracts. mBio. 2015;6(6):e01905–15. doi: 10.1128/mBio.01905-15.

